# Evidence of individual differences in motives for nicotine seeking in classical nicotine self-administration and associated outcomes of varenicline administration

**DOI:** 10.1101/2021.10.05.463198

**Authors:** Vernon Garcia-Rivas, Jean-François Fiancette, Jessica Tostain, Giulia de Maio, Jean-François Wiart, Jean-Michel Gaulier, Véronique Deroche-Gamonet

## Abstract

**Background:** Smokers vary in their motives for tobacco seeking, suggesting that they could benefit from personalized treatments. However, these variations have received little attention in animal models for the study of tobacco dependence. In the most classically used model, ie. intravenous self-administration of nicotine in the rat, seeking behaviour is reinforced by the combination of intravenous nicotine with a discrete stimulus (eg. discrete cue light). In both human and animals, two types of psychopharmacological interactions between nicotine and environmental stimuli have been evidenced. Whether these two types of interactions contribute equally to nicotine seeking in all individuals is unknown.

**Methods:** We combined behavioural pharmacology and clustering analysis. In an outbred male rat population, we tested whether nicotine and the discrete nicotine-associated cue light contributed equally to self-administration in all individuals. Two clusters of rats were identified, in which we further studied the nature of the psychopharmacological interaction between nicotine and the cue, as well as the response to the cessation aid varenicline when nicotine was withdrawn.

**Results:** Notably, withdrawing nicotine produced drastic opposed effects on seeking behavior in the two identified clusters of rats; a 50% increase vs a 18% decrease, respectively. The first cluster of rats sought for the primary reinforcing effects of nicotine and the discrete cue light that has gained nicotine-like secondary reinforcing properties. The second cluster sought nicotine for its ability to enhance the primary reinforcing effects of the discrete cue light. Critically, the approved cessation aid Varenicline counteracted the absence of nicotine in both, but eventually decreasing seeking in the former but increasing it in the latter.

**Conclusions:** Classical rodent models for the study of the reinforcing and addictive effects of nicotine hide individual variations in the psychopharmacological motives supporting seeking behavior. These variations may be a decisive asset for improving their predictive validity in the perspective of precision medicine for smoking cessation.

## Introduction

The main component of tobacco, nicotine, is recognized as one of the most addictive drugs, making smoking cessation difficult, even when 70% of smokers wish to do so (1). From all patients treated with Varenicline (Champix^®^ or Chantix^®^), which is one of the most effective approved pharmacotherapies in supporting smoking cessation (2,3), only 40% remain abstinent at the end of a 12-week-long treatment (4–6). In general, medical treatments are designed and applied as if they would equally benefit to all patients, which is rarely the case. Precision medicine aims at moving from this “one-size-fits-all” approach, by identifying markers of treatment efficacy. Particularly advanced in oncology, precision medicine is limited in psychopathologies such as drug addiction (7).

A major motive for smoking is seeking for nicotine, which is recognized as the main psychoactive compound of tobacco responsible for dependence (8). The paradoxical contrast between the strong addictive profile of tobacco and the relatively weak primary reinforcing effect of nicotine (9,10) has been explained by both clinical and preclinical studies consistently demonstrating that complex interactions between environmental cues and nicotine also play a critical role in promoting and maintaining nicotine seeking (9,11–19). Two major types of interactions between nicotine and environmental cues have been described. Firstly, nicotine seeking can be elicited by external or internal cues after they have acquired nicotine-like effects through Pavlovian association with the primary reinforcing effects of nicotine. Secondly, nicotine can be sought independent of its reinforcing effects, but for its pharmacological ability to enhance the reinforcing value of natural reinforcers (9,12,19–23).

While clinical studies support that the population of smokers is heterogeneous in regards to the breadth of motives that determine the urge to smoke, animal models for the study of tobacco dependence largely ignore this heterogeneity, possibly limiting their predictive validity (24–27). The classical, and most widely used, preclinical model of nicotine intravenous self-administration in the rat involves the contingent delivery of nicotine with a visual stimulus (‘cue’) (28). We and others have shown that this type of visual stimuli is not neutral, but drives self-administration behavior by itself, working as a mild primary reinforcer (22,29–32). Hence, the two types of psychopharmacological interactions between the nicotine and the visual cue can contribute to *nicotine+cue* self-administration. Whether they contribute equally in all rats is unknown. Therefore we explored qualitative differences in the interactions between nicotine and the associated discrete cue as a source of individual variations in the motives for nicotine seeking, as we have proposed before (33). We tested the hypothesis that nicotine and conditioned stimuli may exert a higher control on seeking behavior in some individuals, while others may seek nicotine for its ability to play as a reinforcer-enhancer.

To this end, rats were trained for intravenous nicotine self-administration associated with a discrete visual cue. The respective contribution of nicotine and of the cue to the self-administration behavior was tested by omission of one (Nicotine Omission test) or the other (Cue Omission test). An unbiased clustering analysis was run on variables that accounted for the behavioural effects of the Nicotine and Cue Omission tests. Two clusters were identified with differences in the impact of cue and nicotine omissions on self-administration behavior. They were then characterized for their sensitivity to nicotine reinforcing effects and the nature of the nicotine-cue interaction. Eventually they were compared for the ability of varenicline to reduce seeking for nicotine during a nicotine omission test.

In parallel, controlled experiments were run to confirm that the discrete visual cue was a primary reinforcer whose properties were enhanced by nicotine.

## Methods

### Animals

Male Sprague–Dawley rats (Charles River, France), weighing 280–300 g at the beginning of the experiments, were single housed under a 12 h reverse dark/light cycle. In the animal house, temperature (22 ± 1°C) and humidity (60 ± 5%) were controlled. Rats were habituated to environmental conditions and experimental handling for 15 days before initiation of the experimental procedure. Standard chow food and water were provided *ad libitum*. All procedures involving animal experimentation and experimental protocols were approved by the Animal Care Committee of Bordeaux (CEEA50, N° 50120168-A) and were conducted in accordance with the guidelines of the European Union Directive 2010/63/EU regulating animal research.

### Surgeries

#### Catheterization

A silastic catheter (internal diameter = 0.28 mm; external diameter = 0.61 mm; dead volume = 12μl) was implanted in the right jugular vein under ketamine / xylazine anesthesia. The proximal end reached the right atrium through the right jugular vein, whereas the back-mount passed under the skin and protruded from the mid-scapular region. Rats were given 5-7 days recovery before intravenous self-administration training began.

### Drugs

Ketamine hydrochloride (80 mg/kg) (Imalgène 1000; Rhône Mérieux, Lyon, France) and xylazine hydrochloride (16 mg/kg) (Rompun; Rhône Mérieux, Lyon, France) were mixed with saline and administered intraperitoneally in a volume of 2 ml/kg of body weight. For intravenous self-administration experiments, (−)-nicotine-hydrogen-tartrate (Glentham, UK) was dissolved in sterile 0.9% physiological saline for a final dose of 0.04 mg/kg free base, which was self-administered by the rats via intravenous (i.v.) route in a volume of 40μl per self-infusion. Nicotine solutions with concentrations different to the training dose (0.02mg/kg and 0.06mg/kg free base) were prepared afresh, and used instead of the training dose where indicated. For cue self-administration experiments, a nicotine solution for intraperitoneal (i.p.) administration was prepared by dissolving (−)-nicotine-hydrogen-tartrate (Glentham, UK) in sterile 0.9% physiological saline for a dose of 0.4mg/kg free base, in a volume of 1 ml/kg. All nicotine solutions were adjusted to a pH of 7.

Varenicline or 7,8,9,10-Tetrahydro-6,10-methano-6H-pyrazino[2,3-h] [3]benzazepine tartrate (Tocris, UK) was dissolved in sterile 0.9% physiological saline for a final dose of 1 mg/kg free base. Varenicline was administered i.p. in a volume of 2.5 ml/kg.

### Self-administration

#### Self-administration Apparatus

The self-administration setup consisted in 48 self-administration chambers made of plexiglas and metal (Imetronic, France). Each chamber (40 cm long x 30 cm width x 36 cm high) was located in an opaque sound-attenuating cubicle equipped with an exhaust fan to assure air renewal and mask background noise. Each chamber was equipped with: (a) Photocell beams, located at 1.5 cm from the floor, which allow to record horizontal locomotor activity; (b) Two holes, located at opposite sides of the chamber at 5.5 cm from the grid floor; (c) A common white light (white LED, Seoul Semiconductor, South Korea, 5 Lux), 1.8 cm in diameter, located 8.5 cm above one hole, and commonly designed as cue light; (d) A pump driving a syringe (infusion speed: 20μl / sec) located outside the chamber on the opaque cubicle. Nose-poke visits into holes (see procedures) were used as operant manipulanda to drive and record instrumental responding. Nose-pokes could be reinforced by the cue light only (**Cue Self-administration experiments**), or be reinforced by an i.v. infusion via the jugular catheter through the pump-driven syringe associated or not with the cue light (**Intravenous Self-administration experiments**). Experimental contingencies were controlled and data was collected with a PC-windows-compatible SK_AA software (Imetronic, France).

#### Cue Self-administration Procedures

##### Habituation to nicotine

One week before training for cue self-administration, rats were given daily i.p. injections of nicotine 0.4mg/kg for five days, in order to habituate them to nicotine, to avoid the motor suppressing effects reported after an acute nicotine challenge (34,35).

##### Basic Training Protocol and Pre- and Post-Session injections

At the start of the session, each rat was placed inside one self-administration chamber. Rats were trained for cue self-administration on daily one-hour sessions, running 5 days a week (Monday to Friday). Sessions began two hours after the onset of the dark phase. Nose-poke in the active hole under a fixed ratio 3 schedule of reinforcement (FR3) produced the activation of the cue light located above it (over 4 sec). Nose-pokes at the inactive hole were recorded but had no scheduled consequences. Rats were placed under an FR3 schedule of reinforcement from the first session onwards. Neither food-training, nor FR-1 transition period were used.

During cue self-administration training, rats were given an i.p. injection 5 minutes before the start of the session (“pre-session injection”), and an i.p. injection 3 hours after the end of the session (“post-session injection”). Depending on the experiments and groups, rats received saline as pre-session injection and nicotine (0.4 mg/kg) as post-session injection, while others received the opposite.

##### Test of Varenicline effect on cue self-administration

Varenicline (or its vehicle) was administered i.p. 35 min prior to the cue self-administration session [i.e. 30 min prior to the i.p. (nicotine or saline) pre-session injection], in a volume of 2.5 ml/kg.

#### Intravenous Self-administration Procedures

##### Intravenous Self-administration Basal Protocol

At the start of the session, each rat was placed inside one chamber and connected to the pump-driven syringe (infusion speed: 20μl / sec) through its chronically implanted i.v. catheter. Rats were trained for i.v. *nicotine + cue* or i.v. *saline + cue* self-administration on daily 3-hour sessions, running 5 days a week (Monday to Friday), except for the first session, which took place on a Tuesday. Sessions began two hours after the onset of the dark phase. Nose-poke in the active hole under an FR3 schedule produced the simultaneous activation of the infusion pump (40 μl over 2 seconds) and the cue light located above it (over 4 sec). Nose-pokes at the inactive hole were recorded but had no scheduled consequences, unless otherwise specified. As for cue self-administration, rats were placed under an FR3 schedule of reinforcement from the first session onwards. Neither food-training, nor FR-1 transition period was used. Rats had no limit to the number of self-infusions available. To maintain catheter patency, catheters were flushed with ~10 μl of heparinized saline (30 IU/ml) after each self-administration session, and before the self-administration sessions run on Monday.

During basal nicotine self-administration sessions, two variables were taken into consideration: (1) **the total number of infusions per session** and (2) **a loading index** calculated as the % of infusions achieved after 60 mins. The 60 mins threshold was chosen as it is past the inflection point in the rate of infusions, separating an initial loading phase from a more stable pattern of responding (**FigS1**).

##### Intravenous Self-administration Tests

###### Cue and nicotine Omission tests

To assess responding to nicotine in the absence of the cue, rats performed sessions similar to a standard session, except that nicotine infusions were not paired with the presentation of the contingent cue for the whole duration of the session. Similarly, to assess responding to cue in the absence of nicotine, rats performed a session similar to a standard session, except that the nicotine solution was replaced with 0.9% physiological saline for the whole duration of the session.

For each rat, the effect of cue or nicotine omission on self-administration was evaluated through two variables: **(1)** The *Omission Global Effect (Om-GE)* calculated as the % change in total number of infusions produced by the omission: [*CueOm- or NicOm- GE* = ([total infusions in omission test – total infusions in baseline]/total infusions in baseline) x 100]. This provides quantitative information about the overall effect of the omission test on infusions normally achieved during baseline sessions (**FigS2**); and **(2)** the *Omission Loading Index Effect (Om-LIE)* calculated as the difference in loading index produced by the omission: [*CueOm- or NicOm-LIE*= loading index in omission test – loading index in baseline]. This provides information about the impact of omission on the initial loading of infusions (**FigS2**). For each variable of interest (total infusions or loading index), baseline corresponds to the mean over the two baseline sessions preceding the test (sessions 11 and 12 for CueOm, sessions 16 and 17 for NicOm). For the sake of simplicity, only one baseline (BL) is featured on **FigS2**.

###### Varenicline effect on seeking behavior (nicotine omission)

To assess the effect of varenicline in seeking for the cue in the absence of nicotine, rats performed a “nicotine omission” session preceded by an i.p. injection of varenicline (1mg/kg) 30 minutes before the onset of the session. As a habituation period, rats were handled and received dummy i.p. injections 30 mins before the session during the two days immediately before the varenicline session.

###### Progressive ratio session for nicotine self-administration (cue omission)

To assess the strength of the reinforcing effects of nicotine in the absence of the associated cue, rats performed two consecutive sessions of progressive ratio. These sessions were similar to the “cue omission” session, except that the ratio of responses per nicotine infusion was increased after each infusion according to the following progression: 3, 6, 10, 15, 20, 25, 32, 40, 50, 62, 77, 95, 118, 145, 179, 219 and 268. The maximal number of responses that a rat performed to obtain one infusion (the last ratio completed) is referred to as the breakpoint. The session ceased after either 3 hours or when 17 infusions were reached (the maximal possible number of infusions).

###### Disconnection test

To assess whether responding for the cue was depending on its contingency to nicotine, rats performed five consecutive sessions in which nicotine and cue were no longer contingent. During these sessions nose-pokes at the active hole under FR3 schedule activated the cue, while nose-pokes at the previous inactive hole under FR3 schedule activated the pump-associated syringe, resulting in the delivery of a nicotine infusion. To allow rats to learn that the previous inactive hole was now reinforced by nicotine infusions, nose-pokes at the active holes were not reinforced for the first 30 minutes of the first disconnection session, promoting exploration of the inactive hole.

###### Dose-response sessions for nicotine self-administration (cue omission)

To assess responsiveness to different doses of nicotine, rats performed sessions similar to the “cue omission” session, except that the training nicotine solution (0.04 mg/kg) was replaced by a solution containing either 0.02 mg/kg or 0.06mg/kg nicotine free base. Rats completed at least three consecutive sessions with each new dose.

##### Quantification of plasma nicotine and metabolites

Nicotine (NIC) together with main metabolites, cotinine (COT) and 3 hydroxy cotinine (OHCOT), were determined in rat plasma samples using a liquid chromatography with tandem mass spectrometry detection (LC-MS/MS) method (see **Supplementary Methods** for plasma collection and quantification method).

## Experimental procedures

The timeline of the experiments 1 and 2 is depicted on **FigS3**.

### Experiment 1 – Influence of nicotine on the reinforcing effects of the cue light

After the phase of ‘habituation to nicotine’ (see Methods above), rats (n=45) were trained for cue self-administration for 8 sessions. They received saline i.p. as pre-session injection and nicotine i.p. as post-session injection from sessions 1 to 4. Then the treatment was shifted, i.e. nicotine pre-session and saline post-session, from sessions 5 to 8.

### Experiment 2: Influence of nicotine on the reinforcing effects of the cue light and its modulation by Varenicline

In experiment 1, the influence of nicotine on cue reinforcing effects was tested in rats already experienced with cue self-administration. In experiment 2, we aimed at testing whether nicotine would influence cue self-administration from the first experience with the cue reinforcing effects.

After the phase of ‘habituation to nicotine’ (see Methods above), rats were trained for cue self-administration for 10 sessions. A first group of rats (n=24) received nicotine i.p. as pre-session injection and saline i.p. as post-session injection, from sessions 1 to 9. Based on cue self-administration behavior from sessions 1 to 7, two balanced sub-groups were constituted (n=12 each). On sessions 8 (S8) and 9 (S9), rats were administered with varenicline 1mg/kg i.p. (VAR) or vehicle (Veh) 30 min before the nicotine pre-session injection, according to a latin square design. On session 10, the VehS8-VARS9 sub-group was treated as in sessions 1 to 7 (nicotine pre-session, saline post-session) while the VARS8-VehS9 sub-group was switched to saline pre-session and nicotine post-session.

A second group of rats (n=8) received saline i.p. as pre-session injection and nicotine i.p. as post-session injection, from sessions 1 to 8. On sessions 9 and 10, the reversed treatment was applied, i.e. nicotine pre-session and saline post-session.

### Experiment 3 – Control by nicotine and the visual cue of i.v. nicotine + cue self-administration: Identification and Characterization of Individual Variations

The timeline of experiment 3 is depicted on **Fig1**. 70 rats underwent catheterization surgery, after which they were trained to self-administer *i.v. nicotine + cue (n=62) or i.v. saline+cue (n=8)*, as described above. After an initial training of 12 basal intravenous self-administration sessions, rats underwent a cue omission test (CueOm) on day 13 and a nicotine omission test (NicOm) on day 18. Both tests were separated by basal self-administration sessions.

**Fig1:**
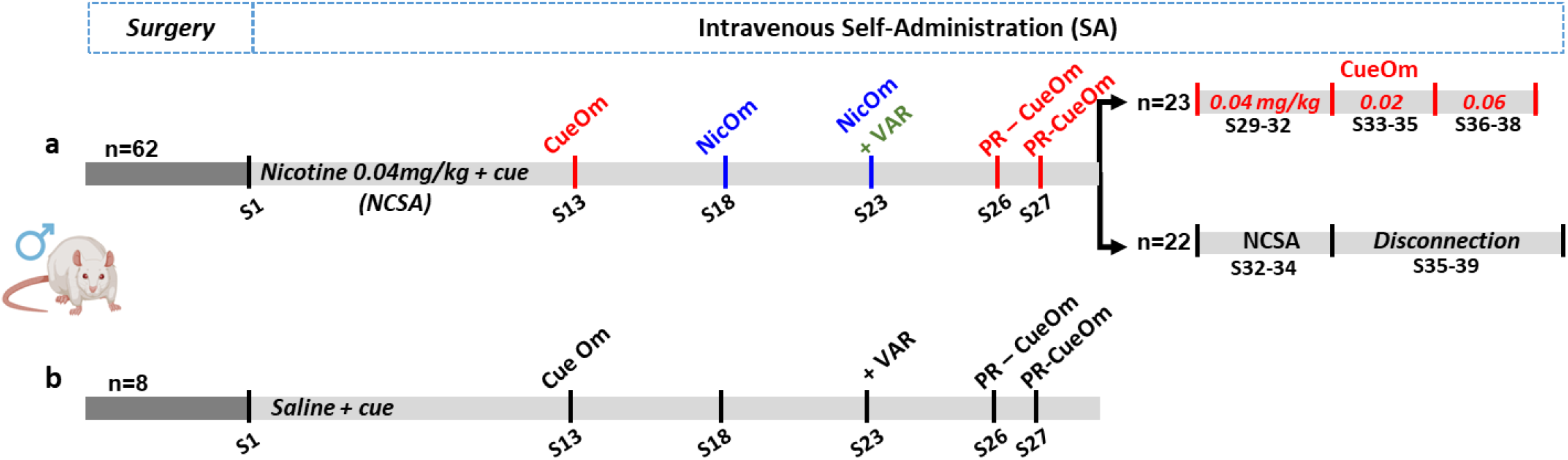
Experimental design of the intravenous self-administration experiment (experiment 3). After for jugular catheterization (*Surgery*), rats were trained for i.v. nicotine+cue (NCSA) (**a**) or i.v. saline+cue self-administration (**b**) at FR3 in 3hrs daily session 5 days per week. The training nicotine dose was 0.04mg/kg/infusion (see details in Methods). After 12 baseline sessions, a series of tests was performed on the whole population up to session 28 or balanced subgroups of NCSA rats starting session 29, to evaluate the contribution of nicotine and the cue to self-administration, the sensitivity to nicotine reinforcing effects and the interactions between nicotine and cue in supporting self-administration behavior: Cue omission on session 13 (CueOm), nicotine omission on session 18 (NicOm), Varenicline effect on a NicOm test (NicOm+VAR) on session 23, Progressive ratio with cue omission (PR-CueOm) on sessions 26 and 27. On sessions 29 to 38, a subgroup of representative NCSA rats was tested in a dose-response for nicotine alone (CueOm), while another representative subgroup was tested in a disconnection test after 7 sessions (28-34) of NCSA baseline. The test consisted in disconnecting nicotine and cue deliveries; the cue remained delivered through active hole visits, while infusions were now delivered in response to inactive hole visits (see Methods for details).

In the *i.v. nicotine+cue* group, an unbiased clustering analysis, based on the four variables of interest CueOm-GE, NicOm-GE, CueOm-LIE and NicOm-LIE, identified two clusters of rats for which nicotine and the cue contributed differently to basal *i.v. nicotine + cue* self-administration. To further characterize these clusters, further behavioral tests were performed in two stages. On the first stage (session 19 to session 28), all rats underwent two tests: (1) *Varenicline effect on nicotine omission test* on session 23, and (2) *Progressive ratio sessions for nicotine self-administration (cue omission)*, on sessions 26 and 27. In all other sessions during this period, rats performed basal self-administration sessions. On the second stage (sessions 29 to 38), 45 rats were selected based on catheter patency and cluster membership, and separated into two different experimental arms: 22 rats underwent the *Disconnection test*, and 23 rats underwent *Dose-response sessions for nicotine self-administration (cue omission)*.

Rats from the *i.v. saline+cue* group went through the same test as the *i.v. nicotine+cue* group up to session 28.

*Experiment 3b*: Twelve rats underwent the same protocol, as the one described for *i.v. nicotine + cue* rats above up to session 18, followed by three additional basal self-administration sessions. Immediately after the end of session 21, 400 μL blood were collected from the catheter for the quantification of plasma nicotine and metabolites (see **Supplementary Methods and Results**).

## Data analyses

### Self-Administration Behavior

Variables of interest in FR self-administration sessions were active and inactive responses, reinforcers earned (infusions or cues), as well as loading index in FR3 sessions and breakpoint in PR sessions for intravenous self-administration. As described above, effects of cue and nicotine omission on nicotine self-administration were evaluated through the *Omission Global Effect (Om-GE)* and the *Omission Loading Index Effect (Om-LIE)*.

### Nicotine and metabolites

In accordance with the literature (36), we used the *main metabolite (cotinine) / parent drug (nicotine) ratio* as an index for metabolism (see **Supplementary Methods**). Different from humans where the two main metabolites, cotinine (COT) and hydroxycotinine (OHCOT) are in a close range, OHCOT levels are low compared to COT in rats, making the OHCOT/COT ratio (nicotine metabolite ratio or NMR) less relevant in rats.

## Statistical analysis

### Self-Administration Behavior

Self-administration behavior was analyzed using one way or repeated measures ANOVA with Time [sessions or time (min) within session], Hole (active vs inactive), treatment (NicOm *vs* NicOm+VAR, VAR *vs* Veh) as within-subject factor, and cluster (cluster A *vs* cluster B) or group (saline pre-/nicotine post-session *vs* nicotine pre-/saline post-session) as between-subject factor. Significant main effects or interactions were explored by pairwise comparisons of means using the Duncan post hoc test.

A factorial analysis was run taking into consideration the four variables obtained from the cue and nicotine omission tests, taken individually for each rat: *CueOm-GE*, *CueOm-LIE*, *NicOm-GE, NicOm-LIE*. An unsupervised *k-Means* cluster analysis was run on the same variables. The *k-Means* cluster analysis was run using a v-fold cross-validation algorithm, which allows for an automatic determination of cluster numbers, without any *a priori* assumptions on the number of clusters to be formed.

All statistical analyses were performed using the STATISTICA 13.3.0 (2017) data analysis software (TIBCO Software Inc, Palo Alto, CA, USA) and TIBCO STATISTICA™ Data Mining system (for unsupervised k-mean clustering). Differences were considered significant at p<0.05. All graphs were done using GraphPad Prism.

## Results

### The cue light has reinforcing effects, which are enhanced by ip nicotine

We evaluated the influence of nicotine pre-session treatment on cue self-administration from the first session (experiment 2) or after 4 sessions under saline pre-session treatment (experiment 1). We show that the discrete cue light associated with nicotine infusions during intravenous nicotine self-administration possesses reinforcing properties by itself and these properties can be enhanced by non-contingent nicotine i.p. administration.

Either in experiment 1 (**FigS4a-c**) or in experiment 2 (**FigS4g-i**), the cue light supported self-administration when the session was preceded by a ip saline injection. In both cases, the number of active responses was higher than the number of inactive responses [Hole effect, F(1,44)=90.28, p<0.0001; **FigS4b** & F(1,7)=9.93, p<0.01; **FigS4h**]. In both experiments, nicotine pre-session enhanced cue self-administration. In experiment 1, the switch from saline to nicotine pre-session was associated with a significant increase in cues earned [TRT effect, F(1,44)=74.2, p<0.0001] (**FigS4a-c**). In experiment 2, the group pre-treated with nicotine from the first session self-administered more than the group pre-treated with saline, over the 7 sessions [Group effect, F(1,30)=13.18, p<0.001] (**FigS4g**); a difference maintained when considering the mean over the last 2 sessions (p<0.01) (**FigS4i**). Notably, nicotine pre-session increased motor activity as compared to saline pre-session, both in experiment 1, when nicotine was substituted to saline pre-session (**FigS4d**) or in experiment 2 when comparing the nicotine pre-session group to the saline pre-session group (**FigS4j**). Enhancement of cue self-administration by nicotine appeared to be independent from altered motor activity, supporting a specific increase of cue reinforcing effects. Not only switching from saline to nicotine pre-session increased nose-pokes specifically in the active hole [TRT x Hole, F(1,44)=93.14, p<0.00001] (**FigS4b-c**), but changes in motor activity [TRT effect, F(1,44)=145.36, p<0.0001] were not related to decreased discrimination between active and inactive holes (**FigS4e**). Similarly, in experiment 2, the nicotine pre-session and the saline pre-session groups differed in motor activity [Group effect, F(1,30)=21.68, p<0.0001] (**FigS4j**), the number of active nose-pokes, but not the number of inactive responses [Group x Hole, F(1,30)=7.96, p<0.01] (**FigS4h**).

Notably, nicotine-induced changes in cue reinforcing effects appeared to be related to the basal reinforcing properties of the cue. The less the cue was reinforcing in baseline conditions (saline pre-session), the more nicotine appeared potent to increase cue self-administration. Indeed, and although r and r^2^ were moderate, increase in cue self-administration by nicotine pre-session was inversely related with baseline cue self-administration under saline pre-session (**FigS4f**).

*Nicotine pre-session* rats from experiment 2 were divided in two equivalent groups (n=12) and tested for the effect of varenicline on sessions 8 and 9, according to a latin square design. Varenicline decreased their cue self-administration behavior (**FigS4k-l**), decreasing specifically active responses (**FigS4l**). Notably, the decrease in cue self-administration produced by varenicline was related to the intensity of cue self-administration in baseline conditions. The higher the self-administration in baseline, the higher the decrease of cues earned by varenicline (**FigS4m**) suggesting that varenicline antagonized the nicotine-induced enhancement of cue reinforcing effects. The group tested with VAR on session 8 and VEH on session 9 was shifted to saline pre-session on session 10 while the first group was maintained on nicotine pre-session. The shift to saline pre-session was associated with a significant decrease in cues earned (**FigS4k middle**). Notably, this decrease was related to the intensity of cue self-administration in baseline conditions. The higher the self-administration in baseline, the higher the decrease of cues earned in response to the switch to saline pre-session (**FigS4n**) further supporting the acute nicotine-induced enhancement of cue reinforcing effects.

### Control by nicotine and the visual cue of *nicotine+cue* self-administration: Identification and Characterization of Individual Variations (Experiment 3)

#### Acquisition of self-administration and contribution of nicotine and cue to self-administration at the population level

##### Saline+cue self-administration

As previously shown (30), rats self-administered *i.v. saline+cue*, as shown by a significant discrimination between active and inactive responses [Hole effect, F(1,7)=35.13, p<0.001], which was stable over sessions [Hole x Session, F(11,77)=1.01, p=0.44], as was the number of reinforcers over the 12 sessions [Session effect, F(11,77)=1.1, p=0.37] (**FigS5a-b**). Cue omission on session 13 induced a significant decrease in self-administration, which resumed on session 14 [Session effect, F(2,14)=3.56, p<0.05], further supporting that the cue was playing as a reinforcer (**FigS5c**).

##### Nicotine+cue self-administration

Over the first 12 sessions, *i.v. nicotine+cue* rats acquired and stabilized self-administration behavior. We observed a progressive increase in self-infusions followed by a plateau around 30 infusions/session [Session effect, F(1,61)=67.78, p<0.0001] (**FigS5d**). Rats nose-poked more in the active hole (whose visit at FR3 triggers delivery of *iv nicotine + cue*) than in the inactive one (whose visits are without scheduled consequence) [Hole effect, F(1,61)=565.8, p<0.0001] and this increased over sessions [Hole x Session, F(11,671)=57.95, p<0.0001] (**FigS5e**). Supporting that *i.v. nicotine+cue* self-administration behavior in our protocol relies on nicotine, the number of infusions earned over a 3hrs session was positively related to the cotinine (main nicotine metabolite) over nicotine ratio (**FigS5f**). Consistent with the literature (37,38), this also supports that high nicotine intake is favored by a fast nicotine metabolism (high cotinine/nicotine ratio).

Not only nicotine, but also the cue contributed to self-administration behavior: both cue omission (CueOm) on session 13 (**Fig2a**) and nicotine omission (NicOm) (**Fig2b**) on session 18 altered the time course of mean infusions as compared to the respective baseline. As compared to baseline, total infusions earned were decreased when behavior was solely reinforced by nicotine (CueOm) [CueOm effect, F(1,61)=256.55, p<0.0001], and tended to be increased when behavior was reinforced solely by the cue (NicOm) [NicOm effect, F(1,61)=2.85, p=0.096] (**Fig2c**). In both cases, the loading index was increased by the omission as compared to the respective baseline [CueOm effect, F(1,61)=41.11, p<0.00001; NicOm effect, F(1,61)=49.35, p<0.00001] (**Fig2d**). CueOm and NicOm had a different impact on self-administration. CueOm and NicOm effects on total infusions as compared to the respective baseline (ie global effect, GE) were different [F(1,61)=83.7, p<0.00001] (**Fig2e**), as were their Effect on Loading Index (LIE) [F(1,61)=7.39, p<0.01] (**Fig2f**).

**Fig2:**
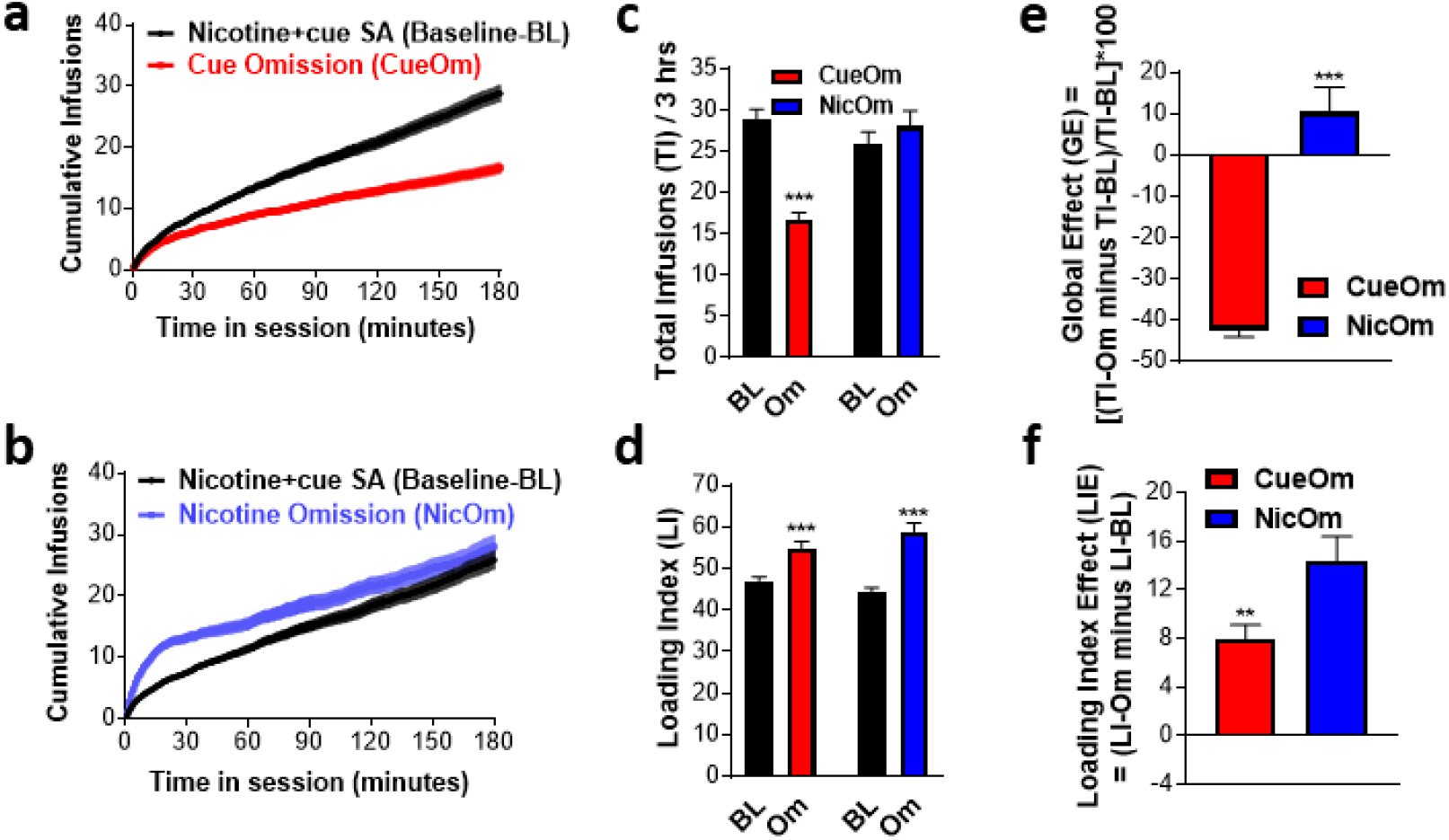
Alteration of *i.v. nicotine+cue* self-administration behavior by cue omission and nicotine omission – population effect. **a**. Cumulative number of self-infusions over the 3 hrs of session. Baseline responding over sessions 11 and 12 is compared to responding over session 13 during which the cue associated with nicotine delivery was omitted (cue omission). **b**. Cumulative number of self-infusions over the 3 hrs of session. Baseline responding over sessions 16 and 17 is compared to responding over session 18 during which nicotine infusions are replaced by saline infusions (nicotine omission). **c**. Total infusions earned during the cue omission and nicotine omission sessions and their respective baseline (BL) sessions. ***p<0.001 as compared to respective BL. d. Loading Index during the cue omission and nicotine omission sessions and their respective baseline (BL) sessions.***p<0.001 as compared to respective BL. Loading Index corresponds to the % of total infusions reached after 60 min. **e**. Global effect (GE) of cue and nicotine omission test corresponding to the mean of individual deltas between infusions on omission test and infusions on corresponding baseline. Cue omission decreased self-administration while nicotine omission increased it. ***p<0.001. **f**. Loading Index Effect (IE) corresponding to the mean of the individual delta between the loading index on omission test and the loading index on corresponding baseline. LIE was higher in response to nicotine omission. **p<0.01. c, e, d, f: Data are expressed as mean±sem.

For recall, to characterize the effect of CueOm and NicOm on self-administration behavior at the individual level, two variables were calculated for each rat [**FigS2**, for simplicity, only one Baseline (BL) is represented]: (**1**)The global effect (GE), which corresponds to the % of change in total infusions as compared to baseline sessions, (**2)** The Loading Index Effect (LIE), which corresponds to the difference in loading Index as compared to baseline sessions. Within each test, we found poor or no relation between the intensity of GE and LIE, supporting that the two variables were not redundant: NicOm-GE and NicOm-LIE were poorly related (r=−0.37, r^2^=0.14, p=0.005), and CueOm-GE and CueOm-LI were unrelated (r=−0.12, r^2^=0.016, p=0.32).

#### Individual variations in the contribution of nicotine and the cue to self-administration

The contribution of cue and nicotine to self-administration were not predictive one of the other. CueOm-GE and NicOm-GE were poorly related (r=0.26, r^2^=0.07, p=0.03), and CuOm-LIE and NicOm-LIE were not related (r=0.005, r^2^=0.0002, p=0.97). We observed large individual variations in the effects of CueOm (**FigS6a-b**) and NicOm (**FigS6c-d**), including variations of opposed signs (**FigS6b-d**). Altogether, this supported that individuals may vary in how nicotine and cue interact to drive nicotine self-administration.

The non-redundancy of the 4 variables of interest was further supported by a factorial analysis showing that the variables were loading on two main different factors (**FigS6e**). Based on these four variables, we run an unbiased k-mean cluster analysis which identified 2 clusters. Cluster A regrouped 34 rats and cluster B regrouped 28 rats. Clusters A and B segregated on the two factors identified through factorial analysis (**FigS6f**), and differed in response to both cue and nicotine omission tests (**Fig3**).

**Fig3:**
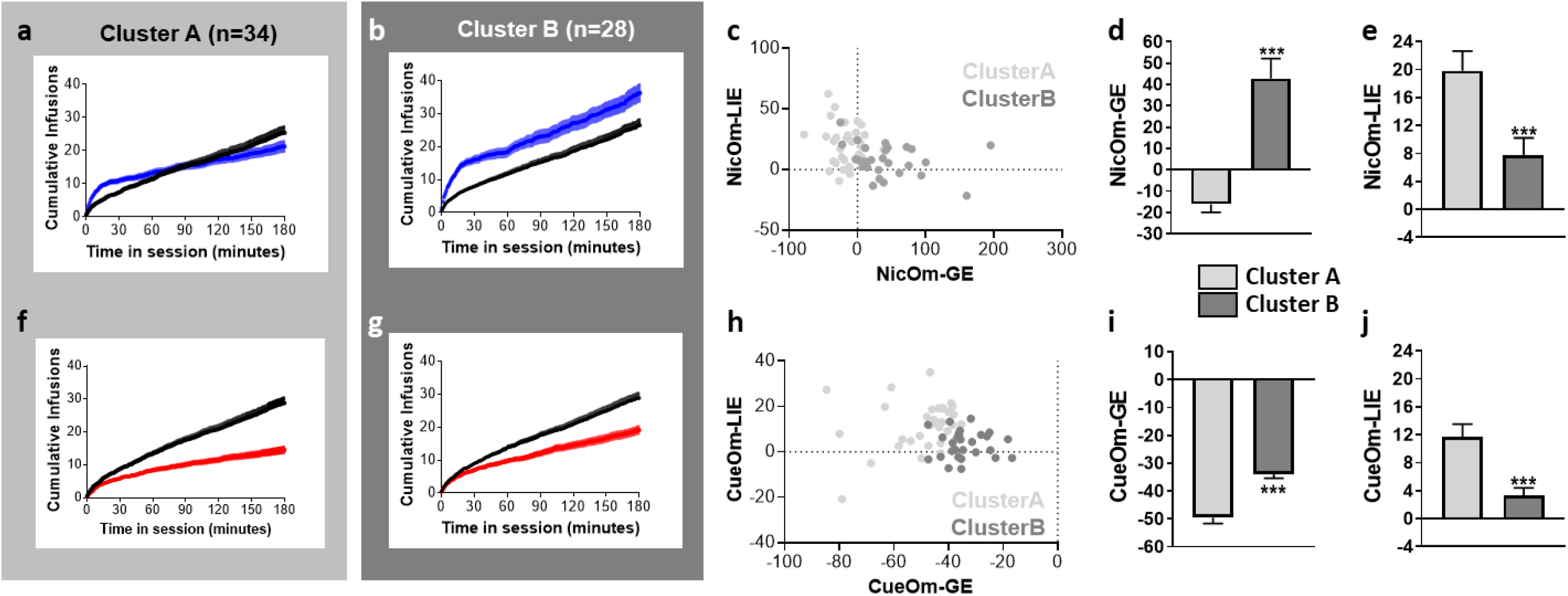
Alteration of *i.v. nicotine+cue* self-administration behavior by cue omission and nicotine omission – Differences between the two identified clusters. **a**. Cumulative number of self-infusions over the 3 hrs session in cluster A (n=34). Baseline responding over sessions 16 and 17 is compared to responding over session 18 during which nicotine infusions were replaced by saline infusions (nicotine omission/NicOm). **b**. Same as **a** for cluster B (n=28). **c**. Distribution of the members of the two clusters in the relationships between NicOm-GE and NicOm-LIE. **d**. Mean NicOm-GE in the two clusters. ***p<0.001. **e**. Mean NicOm-LIE in the two clusters. ***p<0.001. **f**. Cumulative number of self-infusions over the 3 hrs session in cluster A (n=34). Baseline responding over sessions 11 and 12 is compared to responding over session 13 during which the nicotine_associated cue was omitted (Cue omission / CueOm). **g**. Same as **f** for cluster B. **h**. Distribution of the members of the two clusters in the relationships between CueOm-GE and CueOm-LIE. **i**. Mean CueOm-GE in the two clusters. ***p<0.001. **j**. Mean CueOm-LIE in the two clusters. ***p<0.001. d-e, i-j: Data are expressed as mean±sem.

As compared to baseline, nicotine omission decreased total infusions in cluster A, and increased it in cluster B [Cluster x NicOm, F(1,60)=53.92, p<0.00001] (**Fig3a-b**). The two clusters segregated on negative *vs* positive NicOm-GE as well as on low *vs* high NicOm-LIE (**Fig3c**); the two clusters differed indeed for both NicOm-GE [Cluster effect, F(1,60)=38.62, p<0.00001] (**Fig3d**) and NicOm-LIE [Cluster effect, F(1,60)=9.83, p<0.005] (**Fig3e**).

As compared to baseline, cue omission decreased total infusions [CueOm effect, F(1,60)=318, p<0.00001], but less in cluster B than in cluster A [Cluster x CueOm, F(1,60)=9.93, p<0.005] (**Fig3f-g**). The two clusters segregated on higher vs lower CuOm-GE as well as on lower *vs* higher NicOm-LIE (**Fig3h**); the two clusters differed indeed for both CueOm-GE [Cluster effect, F(1,60)=32, p<0.00001] (**Fig3i**) and CueOm-LIE [Cluster effect, F(1,60)=14.08, p<0.0005] (**Fig3j**).

In summary, in cluster A, both nicotine and cue omission produced a decrease in self-administration, supporting that both the cue and nicotine were contributing to self-administration behavior. Differently, in cluster B, the cue alone was able to sustain self-administration. In addition, in this cluster, nicotine alone sustained a higher self-administration behavior than in cluster A. Critically, the two clusters did not differ for acquisition and maintenance of *i.v. nicotine+cue* self-administration (**FigS7a-c**).

The qualitative and quantitative differential effect of CueOm and NicOm in the two clusters (**Fig3**) suggested that their self-administration behavior was supported by a different interaction between nicotine and cue. In cluster A, the two factors, nicotine and cue, would be contributing and necessary to support behavior. Both types of omissions induced an extinction-like effect with both a significant increase in loading index and a decrease in total infusions. Rats from cluster A may be particularly sensitive to the enhancement by nicotine of the reinforcing effects of the cue.

In cluster B, the cue appeared to have acquired secondary reinforcing properties. It was able to support self-administration behavior on its own; increase in loading index by NicOm was associated with an increase in total infusions and a time course of infusions that paralleled the one of baseline (**Fig3b**). Although nicotine alone (CueOm) was associated with an extinction-like profile (increase in loading index and decrease in maximal infusions), it was of a lesser extent as compared to cluster A, suggesting higher reinforcing properties of nicotine in cluster B, consistent with an ability to transfer reinforcing conditioned effects to the cue.

#### Psychopharmacological features of Clusters A and B

To evaluate the primary reinforcing effects of nicotine in the two clusters, we run progressive ratio and FR dose-response tests, in absence of the cue (CueOm). Supporting the results from the CueOm test (**Fig3f-j**) and higher reinforcing effects of nicotine in cluster B, rats from this cluster had a higher breakpoint for nicotine self-administration (**Fig4a-b**). Also their dose-response for nicotine self-administration at FR was shifted upward; they maintain self-administration for a lower nicotine dose (**Fig4c**).

**Fig4:**
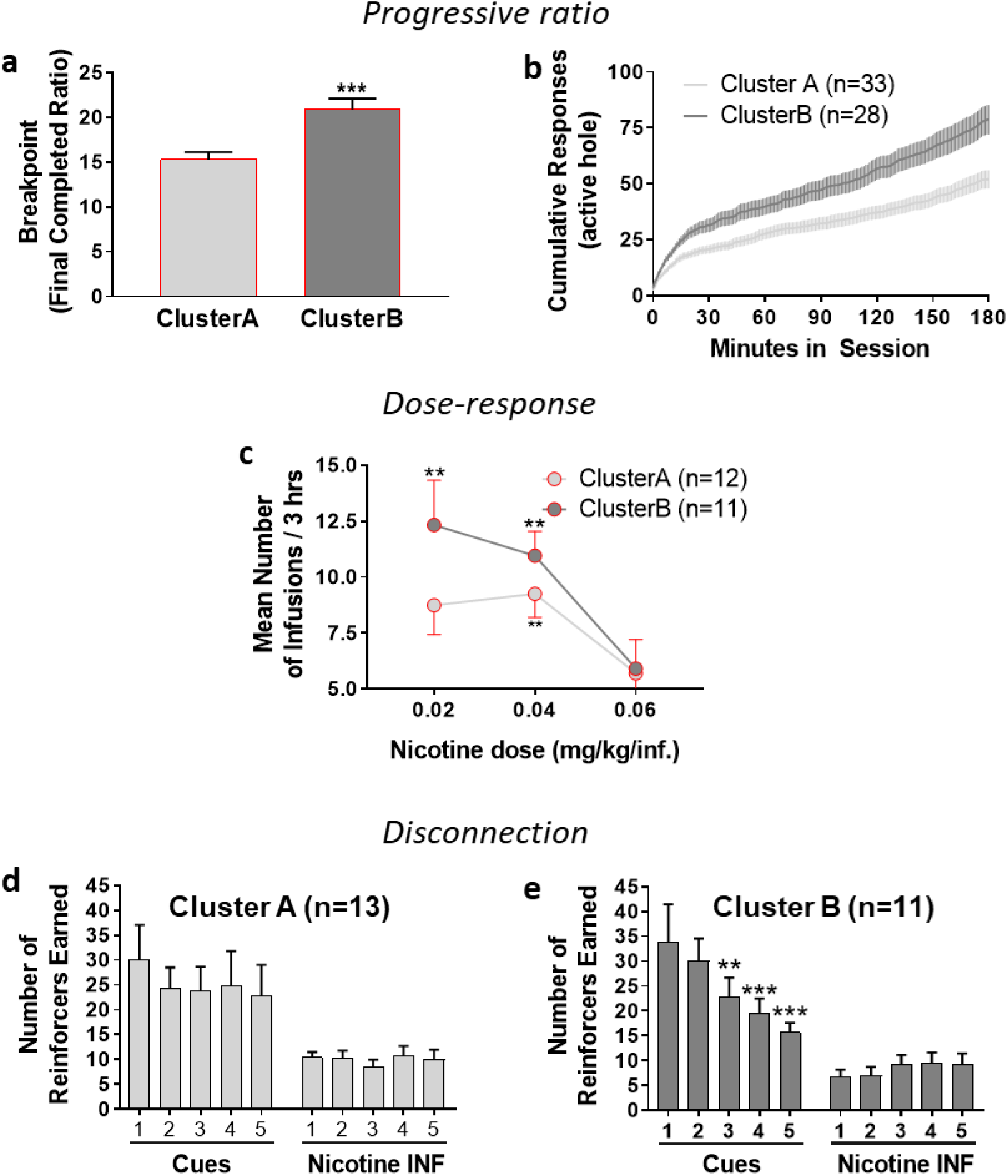
Psychopharmacological characterization of the two identified clusters. The two clusters or representative rats of the two clusters identified based on cue and nicotine omission tests were characterized for their sensitivity to nicotine reinforcing effects and nature of the relationships between nicotine and cue (see Fig1). **a. Mean breakpoint over the two sessions of progressive ratio (S26 and S27)**. Rats were responding for nicotine alone (CueOm). ***p<0.001. **b. Mean cumulative responses in the active hole over the progressive ratio sessions in cluster A and cluster B. c**. Dose-response for nicotine self-administration alone (CueOm) in cluster A and cluster B. **p<0.01 as compared to the respective 0.06 mg/kg dose. **d-e**. Disconnection test in cluster A and B, respectively. For 5 consecutive sessions, nicotine is now delivered by the previous inactive hole, while the cue is delivered by the previous active hole. Disconnection of nicotine and cue did not altr cue resppding in cluster A, but decreased it over sessions in cluster. ***p<0.001, **p<0.01, as compared to the first session. a, c, d-e: Data are expressed as mean±sem.

To test the nature of the interaction between nicotine and cue during self-administration, we run a disconnection test, where cue and nicotine delivery were dissociated and delivered through the active hole and inactive hole, respectively. Rats from the two clusters did not differ for total responding [Cluster effect, F(1,20)=0.09, p=0.76] over the five sessions of test [Cluster x Session, F(4,80)=0.76, p=0.55]. However, they differed in the distribution of responses per hole over sessions [Session x Hole x Cluster, F(4,80)=2.69, p<0.05]. In cluster A, cues earned from responding in the active hole were maintained stable over the five sessions of test, as was the number of nicotine infusions earned from the inactive hole (**Fig4d**). In cluster A, the reinforcing effects of the cue appear to depend on nicotine (**Fig3a**), but do not require nicotine to be contingently delivered (**Fig4d**), consistent with nicotine playing as a reinforcement enhancer even in a non-contingent manner (20,39). In cluster B, cues earned from responding in the active hole decreased over sessions, while the number of nicotine infusions earned from the inactive hole was not significantly affected (**Fig4e**). The reinforcing effects of the cue appeared to be secondary to their contingent association with nicotine delivery and to extinguish if not contingently associated with nicotine, consistent with the cue being conditioned to the primary reinforcing effects of nicotine through Pavlovian conditioning.

#### Varenicline effect on nicotine seeking in Clusters A and B

We evaluated whether Varenicline would differently affect seeking behavior in clusters A and B when nicotine is omitted (**Fig5**).

**Fig5:**
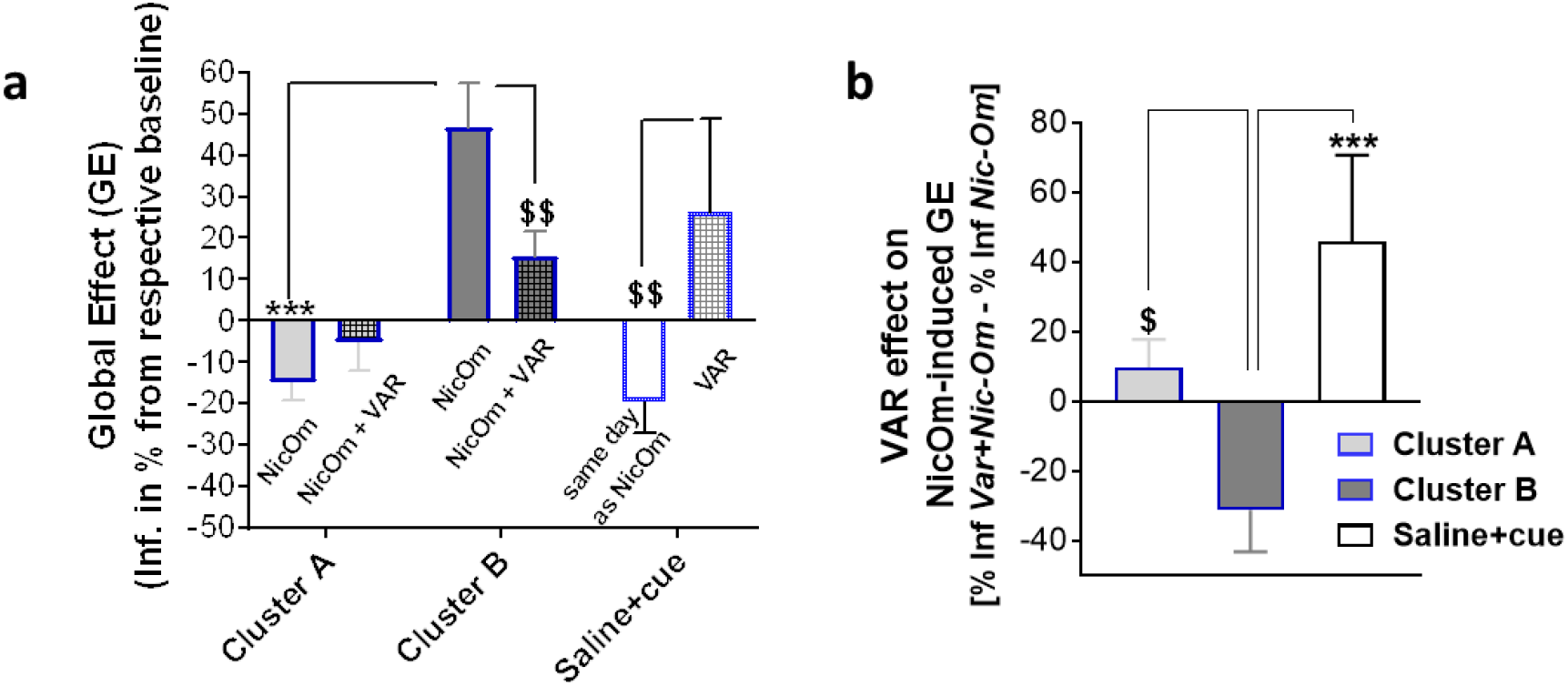
Varenicline effect on responding for the cue (nicotine omission/NicOm) in the two clusters. **a**. Mean Global effect of nicotine omission (NicOm) without (NicOm) or with (NicOm+VAR) Varenicline pre-treatment (VAR). ***p<0.001, $$p<0.01. VAR did not significantly alter cue seeking in cluster A (if anything favored it as compared to simple NicOm) but significantly reduced it in cluster B. In rats self-administering saline+cue, VAR promoted responding for the cue, mimicking nicotine effect. **b**. Compiled VAR effect on nicotine omission global effect in nicotine+cue rats and cue responding in saline+cue rats. $p<0.05, ***p<0.0001. Data are expressed as mean±sem.

Varenicline did not significantly affect self-administration behavior under nicotine omission in cluster A, while it significantly decreased it in cluster B. For recall, Clusters A and B differed significantly when nicotine was omitted (NicOm), cluster A showing a decrease in self-administration and cluster B an increase, as compared to their respective baseline (**Fig5a & Fig3d**). Varenicline reduced the reinforcing effects of the cue (NicOm) in cluster B, but oppositely promoted seeking in cluster A [Cluster x Var effect, F(1,45)=4.06, p<0.05] (**Fig5b**).

Notably, and consistent with a partial agonist effect, varenicline promoted self-administration in the *i.v. saline+cue* control group (**Fig5b**).

## Discussion

In line with clinical studies, animal models for the study of tobacco dependence have consistently demonstrated that various psychopharmacological mechanisms contribute to nicotine seeking (18). Whether all individuals are equal in how these different mechanisms contribute to nicotine seeking is unknown and largely ignored, while smokers may differ in their motives for tobacco seeking [for review (41)]. Based on this, we have recently proposed that the predictive validity of animal models may increase if individual variations are taken into account (33).

In nicotine intravenous self-administration, seeking behaviour is classically reinforced by the combination of intravenous nicotine with a discrete stimulus (eg. a discrete cue light), which is known to play as a mild primary reinforcer by itself. In this context, seeking for nicotine can be driven by the primary reinforcing effects of nicotine in and of itself and by the cue after it has acquired nicotine-like effects through Pavlovian association with nicotine. More recently, it has been shown that nicotine can also be sought after, independent of its reinforcing effects, for its pharmacological ability to enhance the reinforcing value of surrounding natural reinforcers (9,12,19–23). Hence, the two types of psychopharmacological interactions between nicotine and the visual cue can contribute to self-administration. Here, in disentangling the two processes, we show that within an outbred population of male Sprague-Dawley rats, the self-administration behavior is supported by the former in about one half of the population or by the latter in the other half. Varenicline significantly reduced cue seeking in the former but promoted it in the latter.

### Stepwise Omission of the Constituent Components of Classical Nicotine Self-Administration as a Strategy to Reveal Motives for Self-Administration

Since the psychopharmacological interactions between nicotine and environmental cues during classical nicotine self-administration are complex, we developed a behavioral approach that would reveal the independent contributions of both nicotine and cue in the acquired self-administration behavior through omission tests. Previous studies in the past have done similar omission tests separately (20,22,34,42–46), but to the best of our knowledge we are the first to attempt a psychopharmacological profiling of rats based (a) on their individual responses to both omission tests occurring sequentially, and (b) using a clustering method that has been used to identify individual differences without *a priori* assumptions (47). We then developed a series of further tests to better characterize the psychopharmacological profile of these clusters.

### Cluster A: The Reinforcement-Enhancing Effect of Nicotine as a Primary Driver of Nicotine Self-administration

Rats in Cluster A (n=34, 55% of all rats tested) appear driven by the reinforcement-enhancing effects of nicotine on surrounding stimuli, and less by the primary reinforcing actions of nicotine, or cue, by themselves. The defining characteristic of Cluster A is a decrease in overall global responding after both nicotine and cue omission, accompanied with a strong increase in loading index in both occasions (**Fig3e,j**). In other words, during both omission tests rats were approaching an extinction-like profile, in which baseline responding was severely disrupted in the absence of either nicotine or cue. At first glance, this reveals the necessity of both nicotine and cue to be present in order to drive the self-administration behavior in these rats.

Compared to Cluster B rats, rats in Cluster A had lower responsivity to nicotine alone, with a lower breakpoint during progressive ratio (**Fig4a**) and lower dose-responsivity to nicotine in the absence of cue (**Fig4c**). Importantly, Cluster A rats maintained a steady responding for the cue and nicotine even when they were made accessible through different operanda (**Fig4d**), contrasting Cluster B rats, whose response to the cue diminished as its contingency to nicotine was disrupted (**Fig4e**). This non-contingent cue responding is consistent with previous studies showing that the reinforcement-enhancing effects of nicotine do not require previous learning associations, nor contingency with nicotine, to be revealed (20,23,34,48–50). Critically, in replicating the observations reported by Palmatier and colleagues (34), we demonstrate that nicotine can non-contingently enhance responding for the visual cue used in our experimental conditions (**FigS4**), strongly supporting that this psychopharmacological effect by nicotine may be the most important factor in driving nicotine seeking in Cluster A rats.

The reinforcement-enhancing effects on cues exerted by nicotine have been proposed as one of the key mechanisms through which nicotine can be so addictive (19,23), as it can powerfully increase the incentive salience of surrounding stimuli. The behavioral profile of Cluster A evokes the profile of those smokers who consume nicotine for its effect in enhancing the salience of stimuli in their surrounding environment. Even though the reinforcement-enhancing effects of nicotine were first documented in animals (20,22,34,51), there is now substantial evidence of its existence in humans (49,50,52–55). In addition, some studies have proposed a ‘self-medication’ hypothesis of nicotine seeking, in which individuals with socioeconomic or health conditions associated with poor opportunities for reward seek nicotine for its reinforcer-enhancing effects on environmental stimuli (56–59). It is now well documented that sensory anhedonia during nicotine withdrawal can be a strong factor for relapse (60–63). Importantly, in one of our control experiments we demonstrate that nicotine amplifies the responses to a visual cue more in those rats which have a lower baseline responding to the cue in the absence of nicotine (**FigS4f**). It remains unknown whether Cluster A rats had a lower baseline sensitivity to a cue reward in naïve conditions. If that was the case, it could be that they were more sensitive to the reinforcement-enhancing effect of nicotine, making this effect a much more salient event than any reinforcement by nicotine in and of itself. Further studies would be required to explore this possible causality.

### Cluster B: The ‘Classical’ Nicotine-Cue Conditioning as a Primary Driver for Nicotine Self-Administration

Rats in Cluster B (n=28, 45% of all rats tested) appear driven by a combination of the primary reinforcing actions of nicotine and the transformation of the nicotine-paired cue as a conditioned reinforcer, capable of driving self-administration even in the absence of nicotine. The defining characteristic of Cluster B is an increase in overall global responding after nicotine omission, accompanied by only rather small increases in loading index in both cue and nicotine omission (**Fig3e, j**). Removal of the cue decreases their self-administration profile, but not to the extent seen with Cluster A (**Fig3f-g, i**), while removal of nicotine caused an initial sharp increase in responding (**Fig 3b**), followed by a rate of self-infusions not different from their baseline rate. The characteristic response to cue omission could be explained by a heightened sensitivity to the reinforcing actions of nicotine, which, by themselves, appear enough to sustain self-administration. Supporting this, rats in Cluster B have a higher breakpoint for nicotine in the absence of cue during progressive ratio (**Fig4a-b**), and respond more to lower doses of nicotine in the absence of cue (**Fig4c**). When nicotine was omitted, the initial response was a sharp increase in self-infusions, possibly in an attempt to seek for the absent nicotine. The nicotine omission response of this Cluster B is remarkable in that, after this brief initial response, the cue appears to have gained sufficient power as a conditioned reinforcer to drive a rate of self-administration that is not different from their baseline, even in the absence of nicotine. Importantly, when nicotine and cue were made accessible through different operanda, the response for the cue in Cluster B rats diminished over time, while responses for nicotine remained stable (**Fig 4e**). This strongly suggests that for these rats the cue had acquired reinforcement properties due to its contingency with nicotine, which would be extinguished when such contingency was disrupted. This is consistent with previous studies that have shown that nicotine can transform associated cues into conditioned reinforcers, which can sustain self- administration long after nicotine has been discontinued (23,45,64,65).

In human studies, it has been shown that, in some individuals, the environmental stimuli that had become conditioned reinforcers due to their association with nicotine are major sources of craving (66,67), and thus contribute to relapse (68). Some smokers that have been switched to de-nicotinized cigarettes report lower cravings to smoke (65,69,70), suggesting that the conditioned stimuli associated with smoking, such as rolling a cigarette (71), or the oropharyngeal sensations of smoking (72,73), have become strong reinforcers. On the same regard, some smokers report an increase in craving after observing friends smoking, or when visiting the places associated with smoking (17,74–76). Further studies would need to explore whether the observed psychopharmacological profiles in male rats remain the same after protracted nicotine exposure, and whether Cluster B-like rats would be more prone for cue-induced reinstatement.

### Varenicline can have different behavioral outcomes in the absence of nicotine depending on the individual psychopharmacological profile of nicotine seeking

Varenicline is a full agonist at the α7-, and a partial agonist at the α4β2-containing nicotinic cholinergic receptors (77–79). Consistent with its nature as a partial agonist, it can both antagonize nicotine behavioral effects and mimic them in its absence. Our data support this: at a population level, varenicline can moderately enhance cue reinforcing effects in rats self-administering i.v. saline+cue (**Fig5b**), and antagonize nicotine-induced enhancement of cue reinforcing effects (**FigS4k-l**), in accordance with a previous study done in our laboratory (30). Critically, our self-administration data also reveal that the behavioral effects of varenicline in the absence of nicotine can be different, or even opposed, depending on the individual psychopharmacological profile of nicotine-cue interactions: varenicline decreases cue self-administration in cluster B, but slightly increased it in cluster A (**Fig 5a-b**), consistent with the observed effect in reinforcement enhancement (**Fig5b**). Cluster B rats under varenicline strongly diminished the increase in cue responding seen during nicotine omission, suggesting that the partial pharmacological agonism by varenicline was enough to compensate the removal of nicotine, bringing the seeking behavior closer to baseline parameters.

While the therapeutic schedule of varenicline as a tool for tobacco cessation requires daily exposure to the drug in the week leading to a cessation attempt, and a chronic exposure after cessation (80), our study only assessed the effect of an acute exposure to varenicline in the absence of nicotine. The aim of our study was not to test varenicline in a therapeutic perspective, i.e. applied chronically, but as a tool to explore the interactions between nicotine and the cue in the observed subpopulations. Further studies need to assess if prolonged exposure to varenicline affects the psychopharmacological dimensions of nicotine-seeking in the presence and absence of nicotine.

### Individual Differences in Nicotine Seeking: An Opportunity to Improve Preclinical Models of Nicotine Reinforcement

Clinical data strongly support that individuals differ as regards the breadth of motives and mechanisms that determine the urge to smoke [for review (40)], warranting the emergence of research in precision medicine for tobacco dependence. A systematic exploration of individual variations in behavior or pharmacological responses could help improve the translational and predictive value of preclinical models of nicotine reinforcement. Our individual-based approach appears relevant for ‘precision pharmacology’, which at term could be a model for precision medicine in a translational perspective. In the recent years, nicotine metabolism has focused interest as a phenotypic biomarker of smoking heaviness (37) and therapeutic response (81). Fast metabolizers being at risk for heavy smoking (37), and slow metabolizers benefitting from nicotine replacement therapies (NRT) and normal metabolizers from treatments like varenicline (81). Consistently, in rats, the fastest the nicotine clearance, the lowest the nicotine reinforcement threshold and the highest the degree of compensation when decreasing nicotine dose (38), and varenicline appeared more effective at reducing nicotine self-administration in rats with a higher demand for nicotine (82). Interestingly, while in our protocol nicotine demand was positively related to the cotinine/nicotine ratio (**FigS5f**), our two clusters expressed the same self-administration behavior in basal sessions (**FigS7**). It thus remains unlikely that the observed behavioral differences may be primarily due to nicotine metabolism.

Our data support that other markers may predict qualitative variations in mechanisms supporting nicotine seeking. We demonstrated that the same pattern of self-administration behavior can be supported by different psychopharmacological mechanisms, an observation that is in accordance with human evidence (for review, 40). Altogether, the psychopharmacological differences between the two clusters appear independent of extent of nicotine intake during early training. In fact, their psychopharmacological profile only became evident after manipulation of the driving components of seeking behavior: nicotine and its associated cue. Even though the present study was qualitative in nature, it remains a possibility that individuals are sensitive to both psychopharmacological dimensions of nicotine, differing only quantitatively in how much one dimension takes precedence over nicotine seeking during self-administration. The extent to which nicotine and cue control seeking behavior compares between female and male rats, and whether there are sex differences in the proportion of individuals displaying a particular psychopharmacological profile, will deserve attention. Finally, and of special interest to translational approaches, our results open perspective for the study of whether any of these psychopharmacological profiles predict transitioning into compulsive-like nicotine seeking, and whether approved cessation therapies, like varenicline, are more beneficial to individuals fitting a particular profile compared to the other.

Altogether, the evidencing of the individual differences in nicotine seeking reported in this study could help reframe the ongoing discussions about vulnerability to nicotine dependence, explain some of the complexity observed in human and animal studies, as well as providing further insights on why “one-size-fits-all” therapeutic approaches fail to meet the desired clinical efficacy.

## Acknowledgements

This work was supported by La Fondation pour la Recherche Médicale (FRM; grant DPA20140629795 to VD-G), by l’Agence Nationale de la Recherche (ANR; grant ANR-10-EQX-008-1, EquipEx OptoPath to VD-G), by the Aquitaine Region Council (grant 2015-1R30105, PhD fellowship to VG-R), by the LABEX BRAIN (ANR; grant ANR-10-LABX-43, 2019 extension grant fellowship to VG-R), by l’Institut National de la Santé et de la Recherche Médicale (INSERM) and the University of Bordeaux (UB).

We thank S. Laumond, J. Tessaire and the technical staff of the housing and experimental animal facility of the Neurocentre Magendie Inserm U1215. A special thank to Cédric Dupuy for the remarkable care to our rats. We thank Mélodie Nachon-Phanithavong (CHU Lille, Unité Fonctionnelle de Toxicologie, F-59000 Lille, France) for her contribution to analytical analysis.

---

**FigS1:**
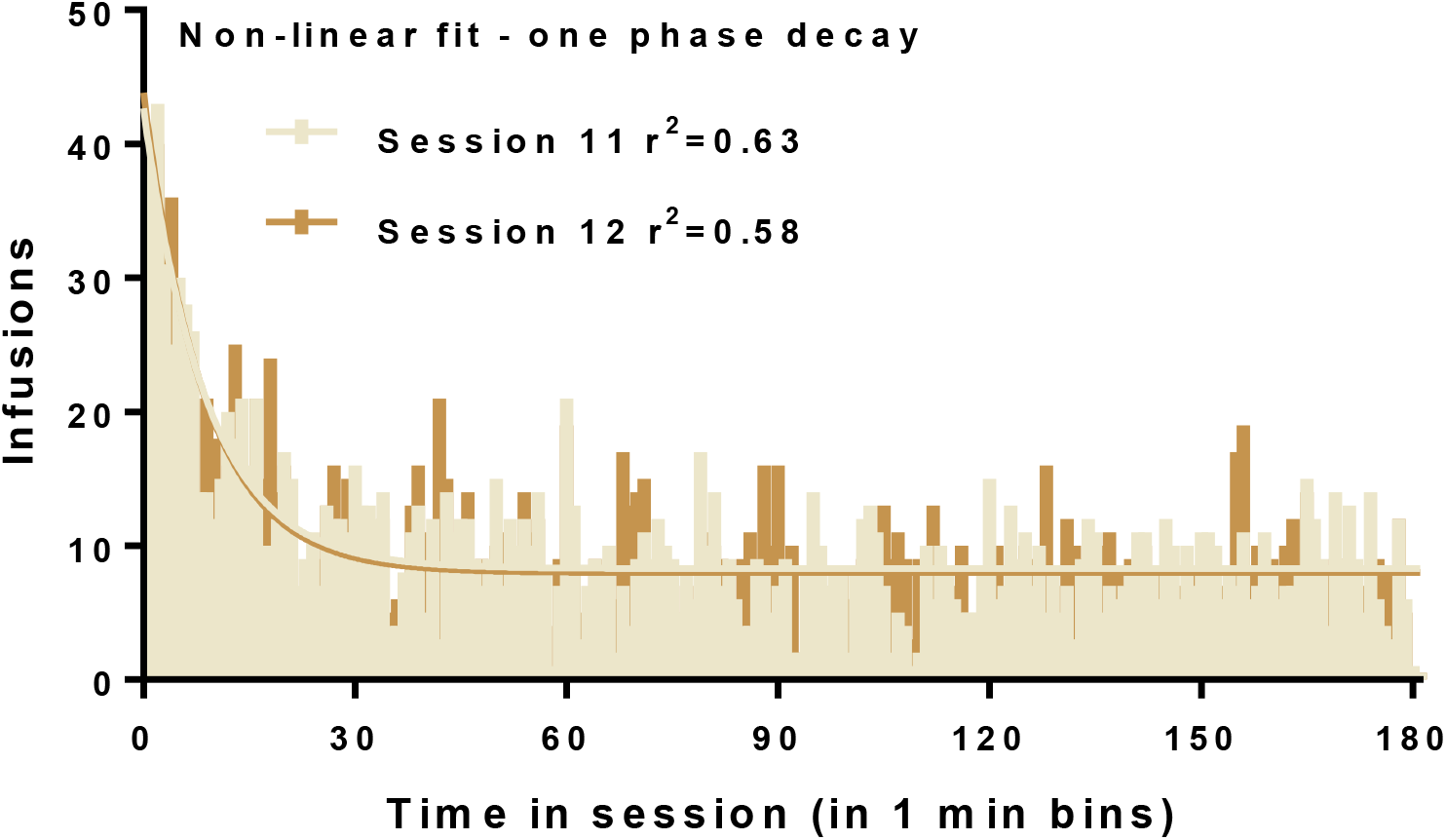
Time course of infusions in rats self-administering i.v. nicotine+cue (experiment 3). Mean infusions per 1 min time bins over the 3hrs of basal self-administration 11 and 12. Time course fitted with a non-linear one phase decay model. After an initial loading phase, intake is regular. Based on this profile, we chose time 60 min to calculate the loading index (% of total infusions at time 60).

**FigS2:**
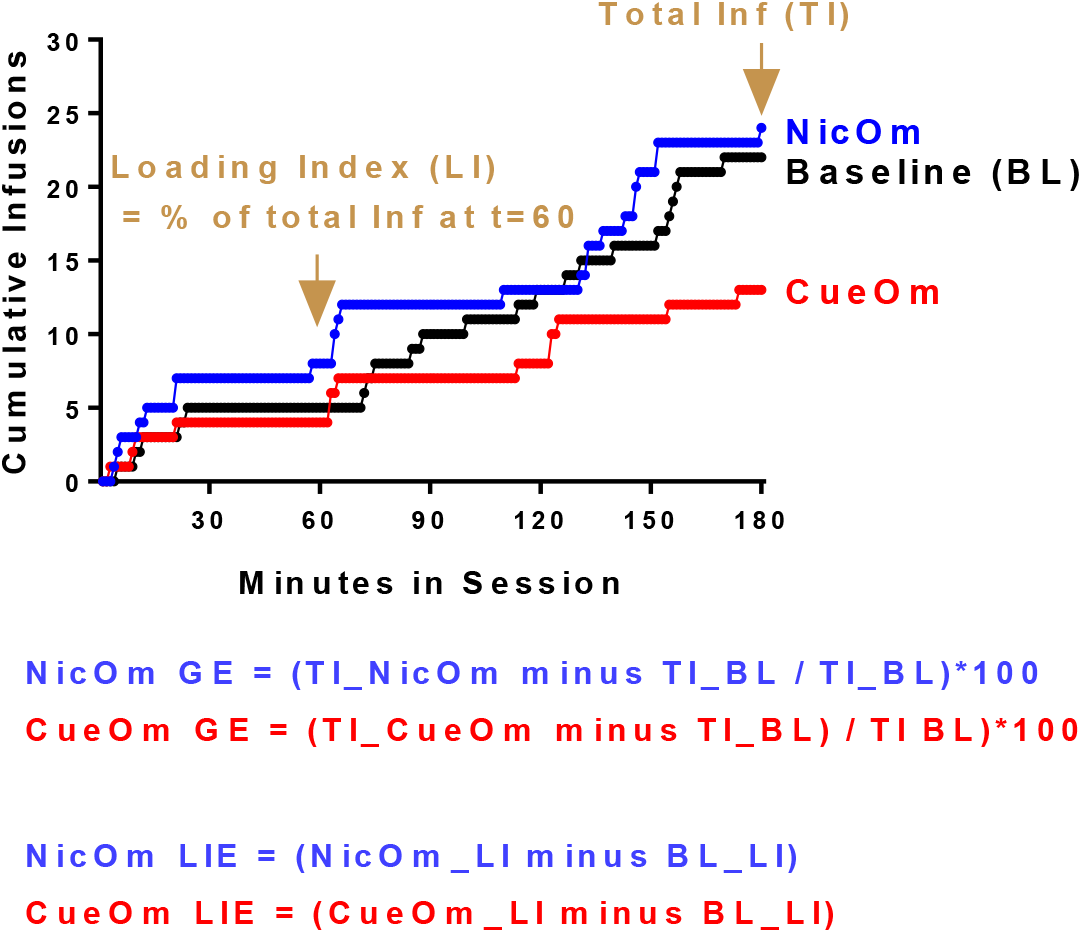
Variables of interest in FR i.v. self-administration sessions and calculation of omission effects. Individual curve of cumulative infusions in cue omission (CueOm), nicotine omission (NicOm) and a baseline (BL) sessions, over 3hrs and per minute. For each rat and session, we considered total infusions (TI) and Loading Index (LI) for the baseline (BL) and respective omission session. Here only one BL is represented for the sake of clarity. Omission Global Effect (Om-GE) corresponds to the impact of omission on total infusions and is expressed as % of baseline. Omission Loading Index Effect (Om-LIE) corresponds to the impact of omission on Loading Index and is expressed as the difference of Loading Index between omission session and respective baseline.

**FigS3:**
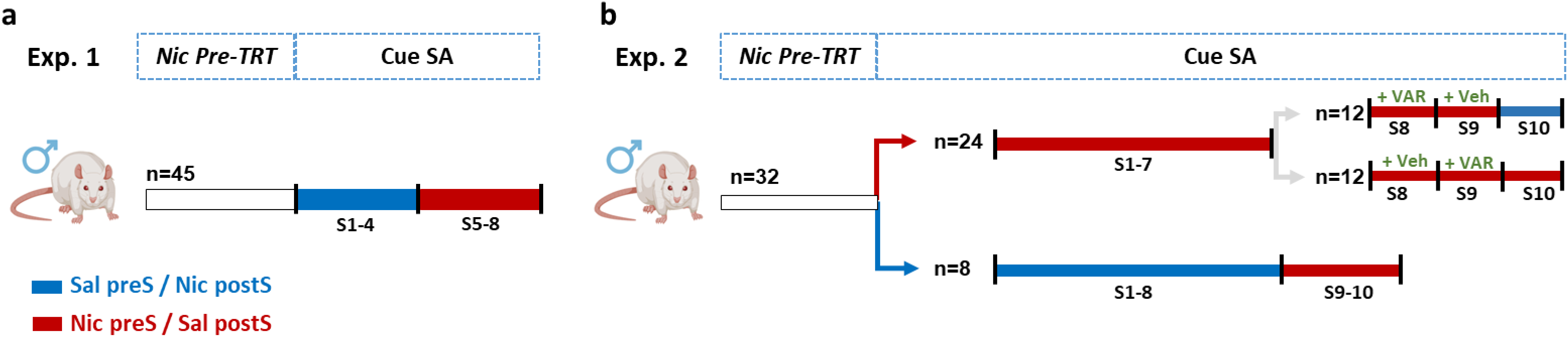
Experimental design of the light cue self-administration experiments testing enhancement of cue reinforcing effects by i.p. nicotine (experiments 1 and 2). In each case, rats were pretreated with nicotine i.p. (0.4 mg/kg, one injection per day for 5 days) to avoid the motor suppressing effects reported after an acute nicotine challenge (34,35). Rats were trained for cue self-administration (Cue SA, 1hr session per day) 5 days per week. Five min and two hours after the session, they were administered i.p. either with saline or nicotine (0.4 mg/kg). **a. In experiment 1 (Exp. 1)**, rats were first trained for 4 sessions with saline pre-session and nicotine post-session (Sal-pre / Nic postS). From sessions 5 to 8, the pre-treatment was switched to nicotine pre-session and saline post-session (Nic preS / Sal postS). **b**. In experiment 2 (Exp.2), the two conditions were run in parallel in two independent groups of rats for 7 to 8 sessions. On sessions 9 and 10, the Sal-preS/Nic postS group was switched to Nic preS/Sal-postS. The Nic preS/Sal postS group was divided in two balanced groups and the effect of varenicline (VAR) was tested according to a latin square design. VAR was administered 30 min prior to the nicotine injection. On session 10, one group was switch to Sal preS/Nic postS while the other came back to baseline condition.

**Figure S4:**
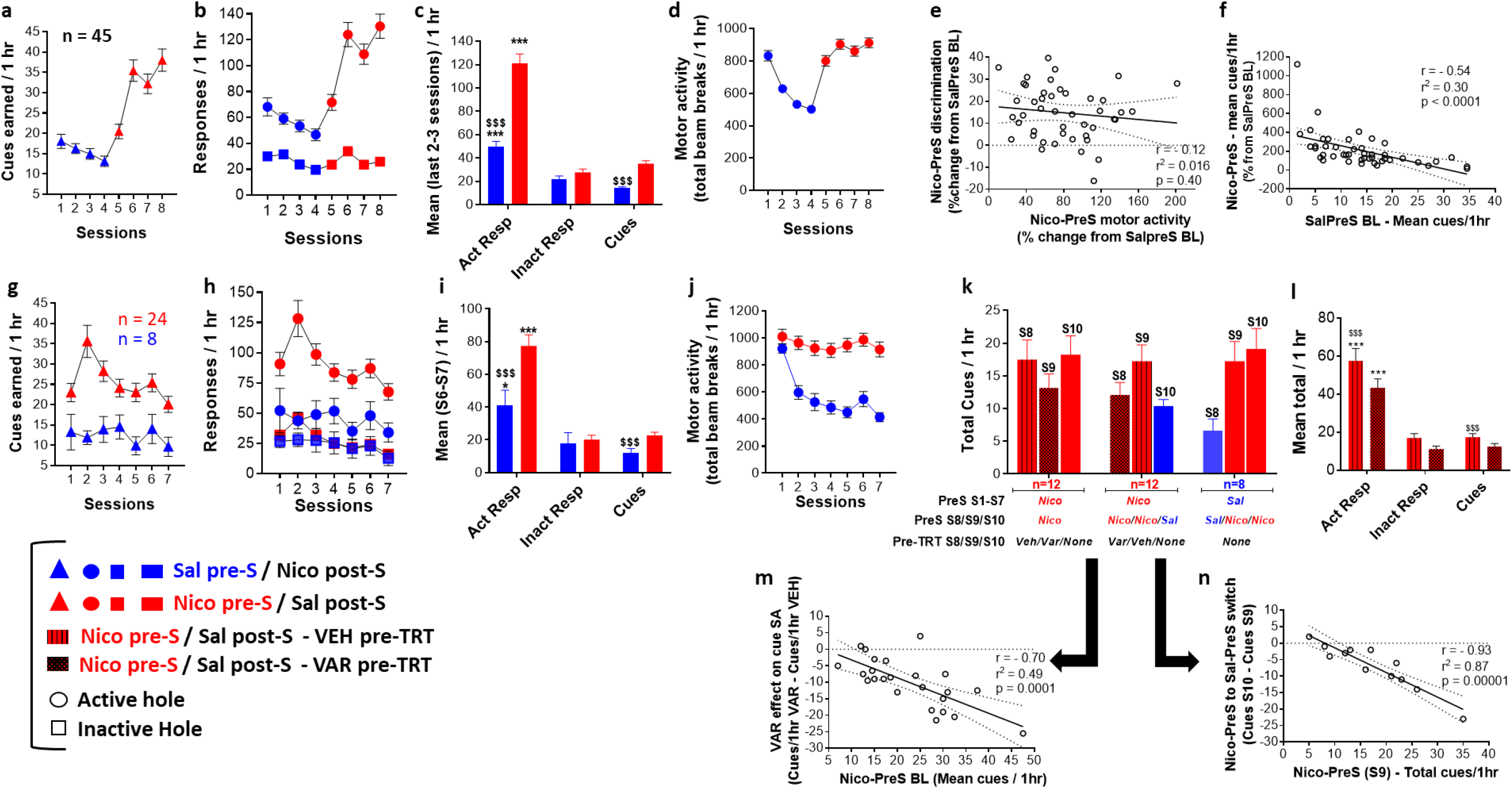
Effect of non-contingent nicotine i.p. on cue self-administration. **a-f** depict results from experiment 1 and **g-n** depict results from experiment 2. **a**. Cues earned per 1hr session. The switch from Sal pre-S to Nic pre-S on session 5 induced an increase in cue self-administration. **b**. Active and inactive responses per 1hr session. The switch o session 5 increased active responses without altering the inactive ones. **c**. Mean responses and cues earned calculated over the last 2 and 3 sessions in Sal pre-S and Nic pre-S conditions, respectively. ***p<0.001 as compared to respective inactive responses, $$$ p<0.001 as compared to Nic pre-S/Sal post-S condition. **d**. Motor activity (as measured by the number of photocell beam breaks) over the 8 cue self-administration sessions. The Sal pre-s / Nic post-S treatment was associated with a progressive decrease in motor activity stabilizing over sessions 3 and 4. The switch to Nic pre-S / Sal post-S induced a stable increase in motor activity. **e**. Changes in motor activity induced by nicotine pre-S was unrelated to changes in hole discrimination. **f**. The increase in cue reinforcing effects in response to the switch to nicotine pre-session treatment was related to the basal reinforcing effects of the cue in the Sal preS condition. **g**. Rats trained for cue self-administration with the Nic pre-S / Sal post-S condition earned more cues than rats trained with the Sal preS / Nic post S condition, from the first session. **h**. Same as g for active and inactive responses. **i**. Mean responses and cues earned calculated over the last 2 in the two experimental groups. ***p<0.001 as compared to respective inactive responses, $$$ p<0.001 as compared to the Nic pre-S/Sal post-S group. **j**. Motor activity (as measured by the number of photocell beam breaks) over the 7 cue self-administration sessions in the two experimental groups. The Nic pre-S / Sal post-S group showed a stable motor activity over the 7 sessions, which was higher than the one of the Sal pre-S / Nic post-S group. In the latter, motor activity decreased over sessions similarly to what was observed in experiment 1 over the first 4 sessions. **k**. Effect of Varenicline and switch to Sal pre-S / Nic post-S or Nic post-S / Sal pre-S on cues earned over 1hr. The Nic pre-S / Sal post-S group was divided in two equivalent groups and tested for the effect of varenicline (VAR) versus Vehicle (VEH) on cue self-administration according to a latin square design run over sessions 8 and 9. When pretreated with VAR prior to nicotine pre-S earned less cues than when pre-treated with VEH. Switch to Sal pre-S on session 10 for one group was associated with a decrease in cues earned. Switch to Nic pre-S / Sal post-S on sessions 9 and 10 in the Sal pre-S / Nic post-S group was associated with an increase in cues earned. **l**. Compiled varenicline effect on active and inactive responses and cues earned, in the 24 tested rats. ***p<0.001 as compared to inactive hole, $$$p<0.001 as compared to VAR pre –TRT. **m**. Effect of varenicline on cues earned was related to basal cues earned under Nic pre-S / Sal post-S pre-treatment. The more the rats self-administered the cue under nicotine effect, the more Var decreased cue self-administration. **n**. Effect of switch to Sal preS / Nic post-S. The more the rats self-administered under nicotine effect, the more the switch to saline pre-session decreased cue self-administration. a-d, g-l: Data are expressed as mean±sem.

**Figure S5:**
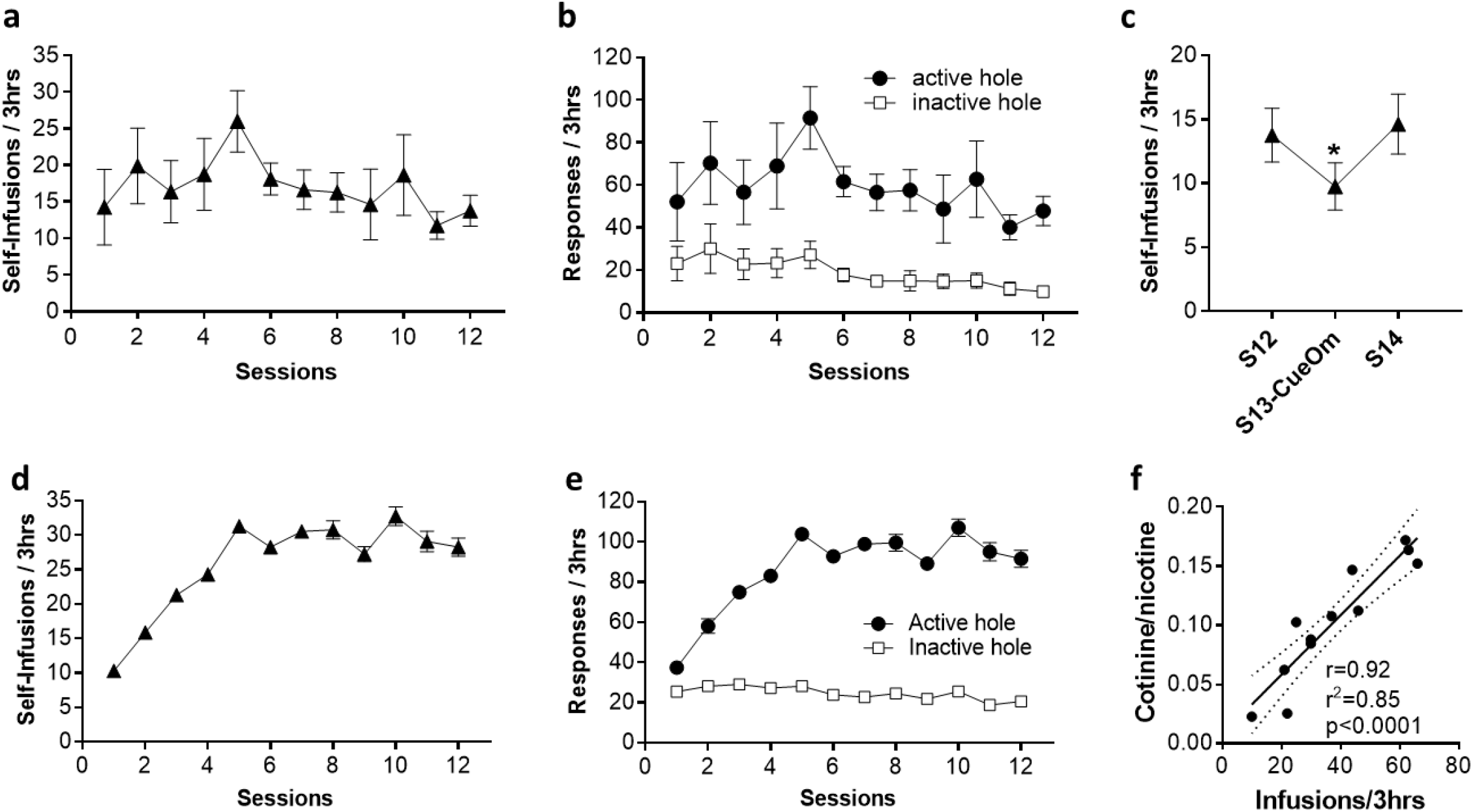
Acquisition of intravenous self-administration behavior in i.v. nicotine+cue and i.v. saline+cue rats and relation of nicotine self-administration with cotinine/nicotine ratio (experiment 3). **a**. Mean self-infusions earned per session by the i.v. saline+cue group. **b**. Active and inactive responses per session by the i.v. saline+cue group. **c**. Effect of cue omission (CueOm) on session 13 in the i.v. saline+cue group. *p<0.05 as compared to session 12 (S12) and session 14 (S14). **d**. Mean self-infusions earned per session by the i.v. nicotine+cue group. **e**. Active and inactive responses per session by the i.v. nicotine+cue group. **a-e**: Data are expressed as mean±sem. **f**. Relationship between nicotine infusions earned over a 3hr-session in i.v. nicotine+cue rats and plasma cotinine/nicotine ratio. The more the rats had self-administered nicotine, the higher the (main metabolite) cotinine over nicotine ratio was high.

**Figure S6:**
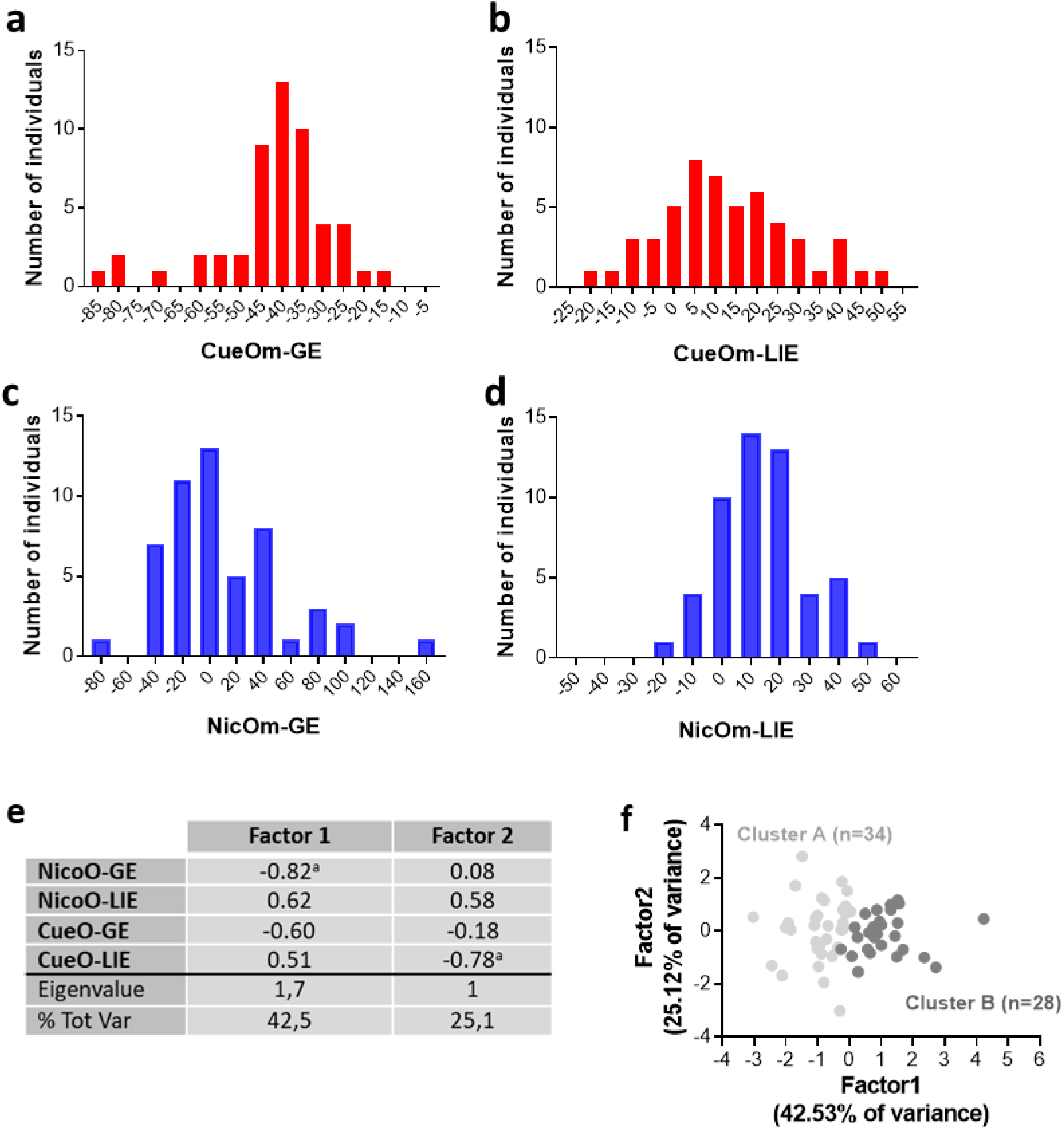
Distribution of the omission scores in the i.v. nicotine+cue group (experiment 3). **a-d**. The scores of the four variables of interest (CueOm-GE, CueOm-LIE, NicOm-GE and NicOm-LIE) were normally distributed with large individual variations including positive and negative scores for the same variable (b-e). **e**. Factorial analysis run on the four variables of interest identified two independent factors. **f**. The scores in the two factors were unrelated at the population level. The two clusters of rats identified through the k means analysis were best segregated through scores on factor 1.

**Figure S7:**
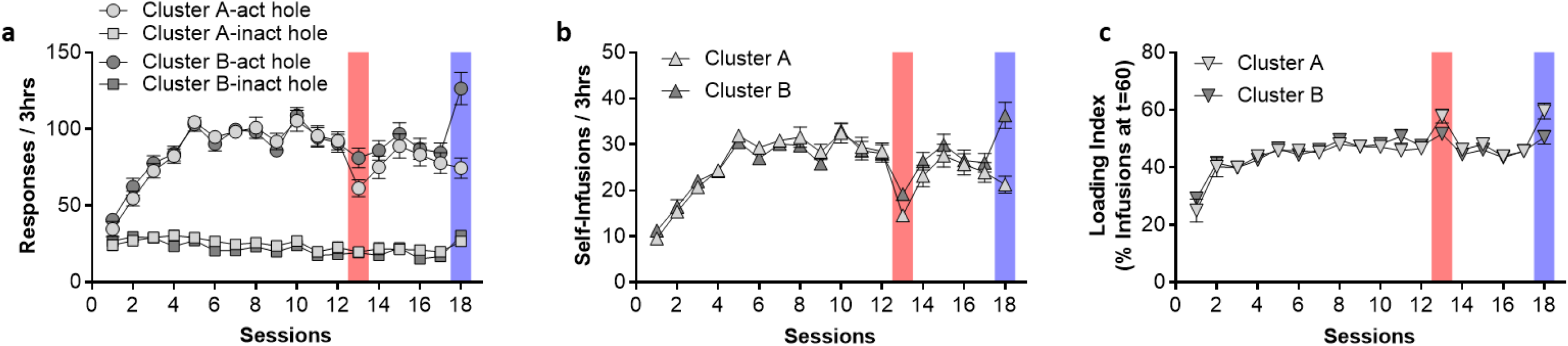
Acquisition of i.v. nicotine+cue self-administration and effect of cue and nicotine omission in the two identified clusters (experiment 3). **a**. Active and inactive responses over the first 18 self-administration sessions in the two clusters. In baseline sessions, rats from the two clusters showed similar levels of responses in both holes. Cue omission on session 12 and nicotine omission on session 18 altered behavior in the two clusters in a different way. **b**. Same as a. for the number of infusions earned per session. **c**. Same as a. and b. for the loading index per session. Data are expressed as mean±sem.

## Supplementary Methods

### Plasma Nicotine and metabolites

#### Plasma collection

400 μL of blood were collected from the catheter immediately after a final basal self-administration session. Blood was put in heparine-containing microtubes (Sarsted 41.1393.005), mixed and placed immediately on ice. Samples were kept on ice until centrifuged (2000 rpm, 10 min, 4°C). Once plasma was separated, 100 μL were carefully pipetted out, placed on 500 μL Eppendorf tubes and stored at −80°C.

#### Quantification of plasma nicotine and metabolites

Nicotine (NIC) together with main metabolites, cotinine (COT) and 3 hydroxy cotinine (OHCOT), were determined in rat plasma samples using a liquid chromatography with tandem mass spectrometry detection (LC-MS/MS) method. Briefly, 100 μL of each plasma sample were mixed with 300 μL of an internal standards (NIC-D_4_, COT-D_3_ and OHCOT-D_3_) mixture in methanol. After centrifugation, 100 μL of 5 mM ammonium formate buffer at pH 3 were added to 100 μL of the obtained supernatant: 10 μL were injected in the chromatographic system. Chromatographic separation was performed using an Acquity™ UPLC HSS C18 (1.8 μm, 2.1×150 mm, Waters) column (Waters) and a gradient of 5 mM ammonium formate / 0.1 % formic acid buffer, and ACN / 0.1 % formic acid as mobile phase. Xevo TQ-S tandem mass spectrometer (Waters) was used for detection after positive electrospray ionization mode in the MRM mode (ESI+) using the following transitions: m/z 163.2 → <u>132</u> and 130 (NIC), m/z 167.1 → <u>136</u> (NIC-D_4_), m/z 177.1 → <u>79.9</u> and 98 (COT), m/z 180.1 → <u>100.9</u> (COT-D_3_), m/z 193.1 → <u>80</u> and 134 (OHCOT) and m/z 196.1 → <u>80</u> (OHCOT-D_3_). In compliance with both the French Analytical Toxicology Society (SFTA) and international recommendations for the validation of new analytical methods (83,84), an additional validation step was performed, which included six independent calibrations conducted on different days and using different rat plasma-free samples. Linearity was determined using linear regression with 1/x weighting for the 3 compounds. The limit of detection (LOD) was defined as the lowest concentration with retention time within ± 0.2 min from the average of all calibrator concentrations and a signal-to-noise ratio of at least three for all selected ion transitions. The lower limit of quantification (LLOQ) was the lowest concentration that could be quantified with acceptable imprecision (CV % ≤ 20 %) and acceptable accuracy (within ± 20 % of the theoretical concentration). Within-day and between-day precision and accuracy were calculated from six repeated analyses of spiked rat plasma samples (at three levels) during one working day, for 6 days.

## Supplementary Results

### Experiment 3b: Plasma nicotine and metabolites

LOD and LLOQ for NIC, COT and OHCOT in plasma samples were 0.5 μg/L. Linear regression with 1/x weighting showed the standard curves (n=6) to be linear from 0.5 to 100 μg/L, with r > 0.999, and observed inter-day CV and bias (n=6) were less than 20%.

Consistent with the literature (36), and different from humans where OHCOT and COT concentrations are in a close range, OHCOT levels were low (15.25 ± 1.2 ng/mL) compared to cotinine (328.7 ± 37 ng/mL) and nicotine levels (3791 ± 564 ng/mL). Hence we used the *main metabolite (COT) / parent drug (NIC) ratio* as an index for metabolism. We found a sustained correlation between this ratio and the total infusions earned (**FigS5f**), while nicotine levels were unrelated to total infusions earned [r=− 0.46, r^2^=0.21, p=0.13]. This supports that variations in nicotine metabolism impact nicotine intake in our basal self-administration procedure in accordance with previous studies (38). Our two clusters did not differ for basal nicotine self-administration behavior; this pleads for no difference in nicotine metabolism between the two clusters.

